# Spatial Probabilistic Mapping of Metabolite Ensembles in Mass Spectrometry Imaging

**DOI:** 10.1101/2021.10.27.466114

**Authors:** Denis Abu Sammour, James L. Cairns, Tobias Boskamp, Tobias Kessler, Carina Ramallo Guevara, Verena Panitz, Ahmed Sadik, Jonas Cordes, Christian Marsching, Mirco Friedrich, Michael Platten, Ivo Wolf, Andreas von Deimling, Christiane A. Opitz, Wolfgang Wick, Carsten Hopf

## Abstract

Mass spectrometry imaging (MSI) vows to enable simultaneous spatially-resolved investigation of hundreds of metabolites in tissue sections, but it still relies on poorly defined ion images for data interpretation. Here, we outline *moleculaR*, a computational framework (https://github.com/CeMOS-Mannheim/moleculaR) that introduces probabilistic mapping and point-for-point statistical testing of metabolites in tissue. It enables collective molecular projections and consequently spatially-resolved investigation of ion milieus, lipid pathways or user-defined biomolecular ensembles within the same image.

---

Mass spectrometry imaging (MSI) has evolved into a label-free core technology for visualization and spatially-resolved analysis of digested proteins, drugs, glycans and metabolites, incl. lipids, in basic, clinical and pharmaceutical research ^1,2^. Despite enormous advances in speed, sensitivity and spatial resolution of MSI instruments and despite a growing number of successful MSI applications, the fundamental concept in MSI data processing, the use of so-called ion images for visualization and molecular analysis, has not changed since the inception of the technology ^3,4^. Ion images, i.e., false color renderings of *m/z* intervals containing an unassigned peak-of-interest (POI), can be prone to technical artifacts ^5^ and user perception-bias ^6^. Moreover, methods for deriving spatial quantitative MSI scores like an overall energy charge score or techniques for spatial probing of global tissue characteristics such as ion milieu, the degree of lipid unsaturation, or even the distribution of entire lipid classes as a function of tissue morphology are lacking.

Here, we report the computational framework *moleculaR* that may replace current ion images by a transformation of MSI data into spatial point patterns (SPP) that, besides a long history in crime pattern analysis, have successfully been used in other biomedical imaging fields like microscopy or magnetic resonance imaging (MRI) ^7–9^. SPPs display points on a map where events, such as presence or absence of metabolites-of-interest (MOI), are likely ^9^. They generally enable new types of arithmetic operations with data, e.g., analysis of localized events and advanced statistical and homogeneity analysis of spatial data ^8^. To this end, *moleculaR* provides the scientific community with solutions for molecular spatial probability mapping and for collective visualization and analysis of molecular ensembles, e.g., alkali metal adducts as ion milieu indicators, energy charge as correlate of metabolic hotspots or entire lipid classes as the basis for the analysis of metabolic pathways.

Ion images currently used in MSI do not represent ion intensity of a single observed POI in a user-unbiased way. They rather neglect mass accuracy and resolving power at the POI *m/z* and use the sum of ion intensities of all peaks present in a user-defined mass range centered on the POI *m/z* instead (**Suppl. Figure 1A**). *m/z* values of observed POI and biologically relevant MOI like the potassium adduct of heme ([Heme+K]^+^), i.e., database entries with corresponding theoretical masses, typically differ. Hence, the molecular identity of POIs is a statistical consideration. Computational frameworks that estimate if POI may correspond to MOI are available, most notably FDR-controlled metabolite annotation provided by platforms like METASPACE (https://metaspace2020.eu)^10^. To complement this POI-centric analytical perspective (is POI=MOI?), we introduce an MOI-centric biomedical perspective that systematically analyzes and visualizes if biologically relevant MOIs have a statistically validated spatial presence across tissue morphologies. To this end, we propose molecular probability maps (MPMs) for rigorous user-independent spatial statistical testing that are based on the assumption that for any given MOI spatial autocorrelations exist that mirror the biological interaction between neighboring tissue morphologies ^11^. Instead of estimating this correlation intensity globally, our approach localizes areas of heightened “activity” in terms of points proximity and signal intensity. We propose that MPMs shall replace or complement ion images for the spatial analysis of MOI.

To generate MPMs, matrix-assisted laser desorption/ionization (MALDI) MSI raw data of any given MOI *m/z* is first transformed into an SPP representation, a well-known basis for statistical data interpretation in other fields of biomedical imaging ^8^. This transformation features data-rather than user-defined mass windows and Gaussian-weighted ion intensities of observed POIs that are based on estimates of the mass resolving power at the MOI’s *m/z* (**Fig. 1A** and **Suppl. Fig. 1B**). Next, to evaluate whether or not point intensities and spatial clustering of the MOI in the tissue section are statistically significant, a complete spatial randomness (CSR) model of that MOI is created by random spatial permutations of MOI points (see Methods), which is then used as the spatial null hypothesis (**Fig. 1B**). Applying kernel density estimation via an isotropic Gaussian with an POI-specific bandwidth estimation (**Suppl. Fig. 2**), the intensity distribution of the CSR density image is expected and observed to converge towards a normal distribution (**Suppl. Fig. 3**), which then forms the basis for inferring intensity cutoffs beyond which the intensities of MOI’s density image are unlikely to occur if generated by a random spatial process (see Methods). MPMs are then defined as the composite representation of MOI spatial distribution on a rastered grid with data-dependent Gaussian weighted intensities (according to the scheme of **Fig. 1A**) with analyte hotspots and/or analyte coldspots superimposed as contours that identify areas of MOI significant abundance and deficiency, respectively (**Fig. 1B**). As exemplified for the sphingomyelin SM(d34:2)+H^+^ (0.1 FDR), MPMs are rather robust against different types of computationally added noise, as evidenced by Dice similarity coefficients of 0.80, 0.97 and 0.98 for comparisons of MPMs based on raw data versus added Gaussian noise, spiked artifacts (isolated very high-intensity pixels) or interfering signals placed in the proximity of the MOI, respectively (**Fig. 1C**). Applying the same testing procedure to 142 MPMs of MOIs (METASPACE-verified at ≤ 0.2 FDR) in positive ion mode revealed median Dice similarity coefficients of 0.76, 0.96 and 0.94 for these three types of added noise, respectively (**Suppl. Fig. 4**). In a neurooncology example, MPMs of two exemplary MOIs, the sphingomyelin SM(d36:4)+H^+^ (0.10 FDR) and the phosphatidylserine PS(36:1)-H^-^ (0.05 FDR), demonstrate how spatial probabilistic mapping of analytes aids in outlining the significant presence or absence of analytes relative to vital tumor regions, as inferred from a neuropathologist’s annotation of a fresh-frozen tissue section of isocitrate dehydrogenase (IDH) wild-type glioblastoma (GB) (**Fig. 1DE**). One important application of MPMs is the spatially-aware analysis of differential distribution of drugs or metabolites between test tissues, e.g. those dosed with drugs or carrying mutations, and corresponding reference tissues. Here, for instance, MPMs map statistically-validated significant localization of immunosuppression-associated tryptophan in IDH-mutant-compared to IDH wild-type glioblastoma ^12^, thereby enabling true spatial probabilistic cross-tissue comparisons (**Fig. 1F**). In such scenarios, intensity weights in the CSR model are sampled from the intensities of the corresponding reference tissue, thus enabling probabilistic cross-tissue comparisons.

**Figure 1.**
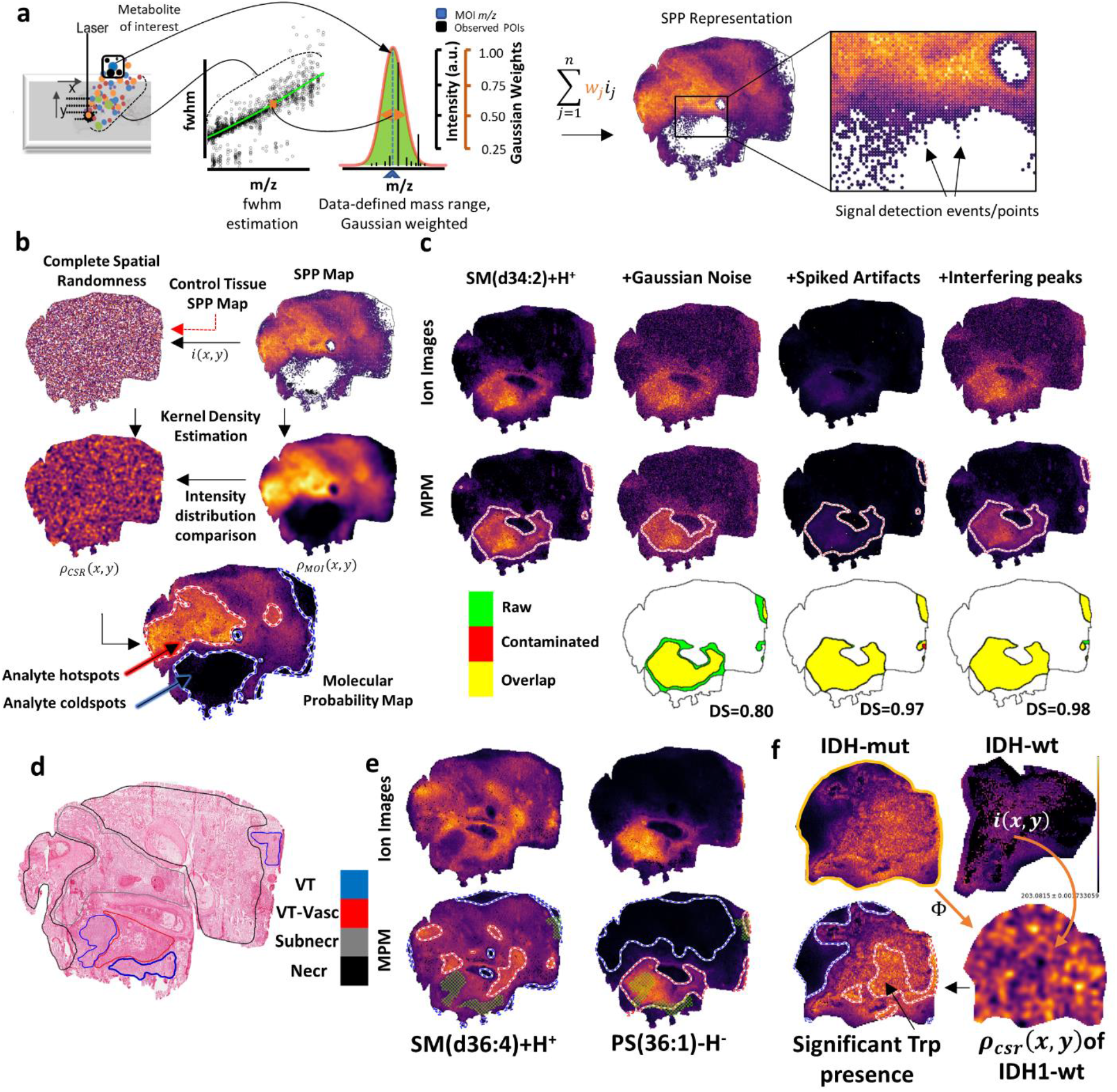
Metabolite Probability Maps (MPM) for spatial probabilistic mapping of metabolites in MALDI-MSI. **a)** SPP representation of metabolite-of-interest (MOI). Full-width-at-half-maximum (FWHM) values are computed for all peaks of a user-selected full mass spectrum and curve-fitted to describe FWHM as a function of *m/z*. For any MOI *m/z* (dashed blue line), a Gaussian envelop is computed with *σ* equaling the estimated FWHM at MOI *m/z*. The observed nearby peaks-of-interest (POIs) are Gaussian-weighted, thus down-weighting proximal background signals. **b)** MPM computational workflow. A corresponding CSR model is created for each MOI’s SPP with equal spatial point density. Kernel density is estimated for both sides; the resulting spatial density functions, *ρ_MOI_*(*x, y*) and *ρ_CSR_*(*x, y*), are compared to estimate areas of significant abundance (MOI “hotspots”; red/white contours) or deficiency (MOI “coldspots”; blue/white contours). **c)** MPMs (middle row) but not ion images (upper row) of exemplary sphingomyelin SM(d34:2)+H^+^ (0.1 FDR) are robust against noise and common signal artifacts: random Gaussian noise (second column), presence of abnormally-high-intensity peak artefacts (third column) and added overlapping peaks 2*σ* away from MOI *m/z* (fourth column). MPM hotspots in raw, noise-free (green) and contaminated data (red) show high overlap (yellow) for all artefacts as judged by their Dice similarities (DS). **d)** H&E-stained image of a human glioblastoma (GB) tissue section (VT: vital tumor; VT-Vasc: vascularized vital tumor; Subnecr: pre-necrotic; Necr: necrotic). **e)** Comparison of ion images and corresponding MPMs of SM(d36:4)-H^-^ and PS(36:1)-H^-^ (≤0.10 FDR) relative to VT regions (green mesh). **f)** MPMs enables spatially-aware cross-tissue comparison of tryptophan [Trp-H]^-^ in IDH-mutant GB as test tissue compared to IDH-wild type GB as reference tissue, where the CSR model of the test tissue is inferred from signal intensities of the reference tissue.

Even more far-reaching than single molecule MPMs, data-integrating probability maps of entire lipid classes in SwissLipids (https://www.swisslipids.org)^13^, of large groups of molecules such as potassium adducts of lipids ^14,15^ or of any other user-defined set of metabolites, so called collective-projection probability maps (CPPM), pave the way for visualization and exploration of integrated MSI data: Here, molecular ensembles are transformed to their respective SPP representations, then collectively projected into a single image space (**Fig. 2A**), and finally subjected to spatial probabilistic mapping into CPPMs (**Fig. 2BCD**). Importantly, this computational framework permits spatial evaluation of composite numeric scores obtained by applying basic arithmetic operations on spatial point patterns of multiple MOIs, for example calculating the spatial abundance of adenine nucleotides [ATP-H]^-^, [ADP-H]^-^ and [AMP-H]^-^ individually relative to their collective sum and a more complex evaluation of adenylate energy charge ^16^ and adenylate kinase mass action ratio ^17^ (**Fig. 2B**). Our data suggests that the latter two scores be indicative of viable tumor (VT), suggesting that CPPMs of molecular ensembles as innovative use of MSI data can provide insights into spatially resolved pathophysiology that would not be possible by single molecule ion images.

**Figure 2.**
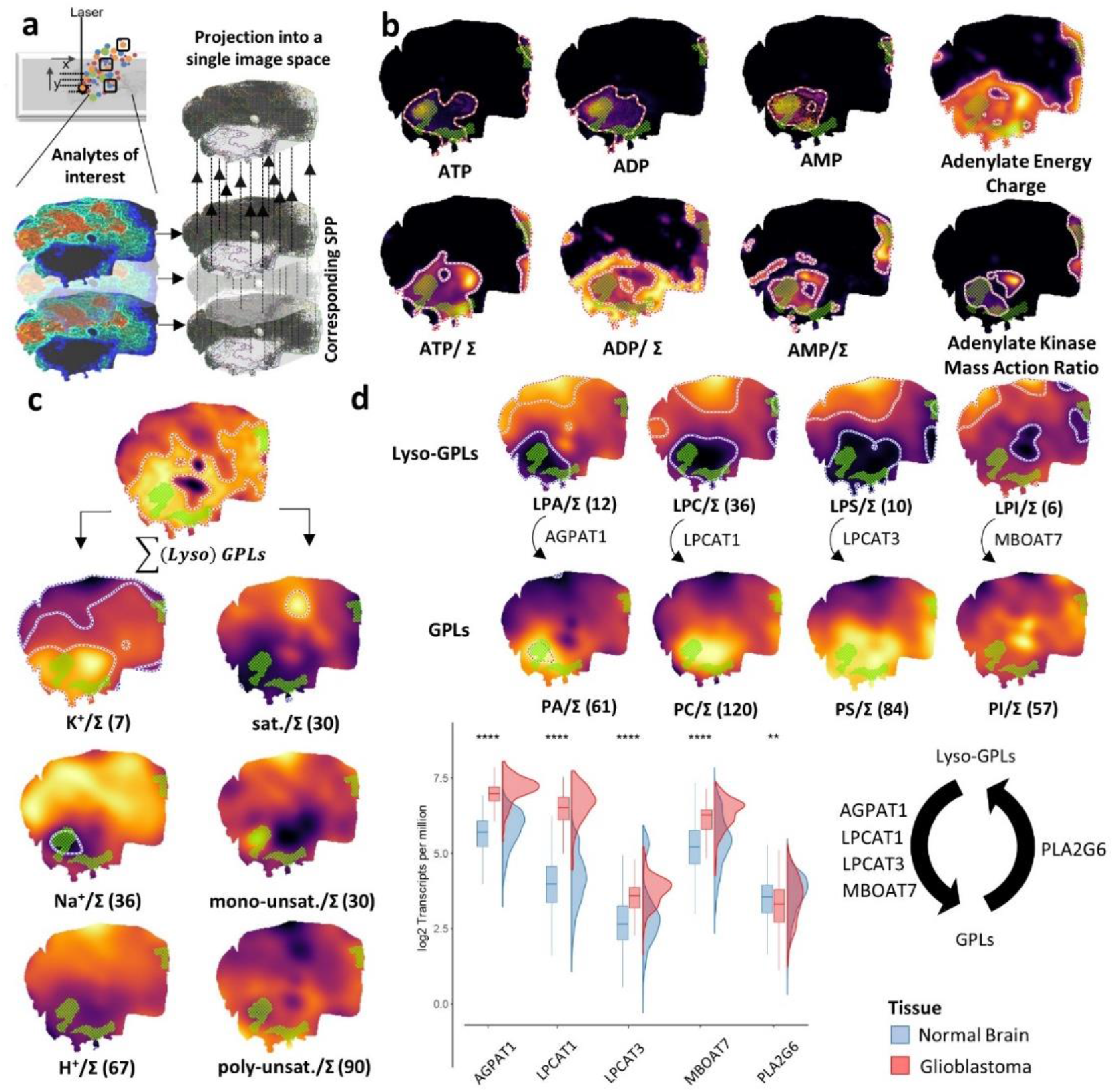
Collective projection probability maps (CPPMs) of molecular ensembles representing scores for energy metabolism, ion milieu, degree of lipid unsaturation, or (lyso)glycerophospholipid remodeling. **a)** SPPs for a cohort of MOIs are collectively projected into the same tissue space. MPMs are then computed from these collective SPPs. **b**) CPPMs enable basic arithmetic manipulations on SPPs of multiple MOIs. Green mesh indicates co-registered vital tumor regions (**Fig. 1d**). MPMs of [ATP-H]^-^, [ADP-H]^-^ and [AMP-H]^-^ (≤0.2 FDR; upper row) compared to their sum-normalized CPPMs (bottom row; Σ = [ATP-H]^-^ + [ADP-H]^-^ + [AMP-H]^-^). CPPMs also enable complex spatial quantitative scores such as adenylate energy charge (([ATP-H]^-^+0.5 [ADP-H]^-^)/([ATP-H]^-^+[ADP-H]^-^ +[AMP-H]^-^); top right) and adenylate kinase mass action ratio ([ATP-H]^-^[AMP-H]^-^/[ADP-H]^-**2**^; bottom right). **c**) Analysis of the tissue’s alkali ion milieu and lipid (un)saturation. Numbers in parenthesis = METASPACE-verified lipids at ≤ 0.2 FDR. left column: CPPMs of all detected protonated or alkali metal adducts of (lyso-)GPLs (PC, LPC, PE, LPE, PS, LPS, PI, LPI) relative to the overall sum Σ(lyso)GPLs. right column: CPPMs showing lipid (un)saturation relative to Σ(lyso)GPLs (sat: saturated; unsat: unsaturated). **d**) CPPMs enable spatial investigation of glycerol-phospholipid remodeling (Lands’ cycle) in GB by visualizing structurally similar lipids (≤ 0.5 FDR) within the same image space. Upper panel: Lyso- and non-lyso-GPL pairs are normalized to their sum (ex. for LPC and PC, Σ represents the sum of all LPC and PC lipids). Lower panel: Rainfall plot showing boxplot and density curves representations of the expression of select Lands’ cycle enzymes in normal brain (blue; GTEx data) and GB (red; TCGA data) both represented as log2 transcripts per million (Wilcoxon rank-sum test, ** P < 0.01, **** P < 0.0001). **Suppl. Fig. 8** illustrates corresponding data for a serial section.

Analytical considerations such as region-specific ion suppression notwithstanding, we reasoned that CPPMs of all potassium or sodium adducts of lipids in the SwissLipids database could serve as indicator of the ion milieu in a cancer tissue sample. Analogously, it has been noted that sodium MRI can serve as an indicator of vital tumor *in-vivo* ^18^: Na^+^/K^+^-ATPase maintains high overall potassium and low tissue sodium concentrations in viable cells, and higher cellularity corresponds to a lower tissue sodium concentration ^18^. In contrast, highly abundant Na^+^-adducts colocalize with necrotic tissue in xenografts of five different tumor cell lines ^15^. Similarly, the projected molecular ensemble (=CPPM) of potassium adducts of all (lyso-) glycerophospholipids (GPLs) was more abundant in vital tumor and vascularized areas, whereas the CPPMs of projected sodium adducts was more pronounced in necrotic tissue showing significant absence (analyte coldspot) in vital tumor (**Fig. 2C**).

Similarly, viable tumor displayed reduced GPLs with short fatty acids (< 34 C-atoms per two fatty acids) (**Suppl. Fig. 5)** or with saturated fatty acids (**Fig. 2C**), but GPLs with 34 to ≤ 40 C-atoms were enriched there (**Suppl. Fig. 5)**. CPPM-based mapping of saturated GPLs indicated derichment in VT areas, but enrichment of mono- and poly-unsaturated GPLs was not as prominent as suggested based on few selected ion images (**Fig. 2C**) ^19^. Finally, collective projections of molecular ensembles support initial surveys of entire molecular pathways. For instance, >400 lipids involved in GPL biosynthesis and remodeling can be interpreted in a single pathway overview (**Suppl. Fig. 6 and 7**). Interestingly, CPPMs of PC, PA, PS but less so of PI and their corresponding lyso-GPL derivatives suggest alterations in the Lands’ cycle of phospholipid remodeling in GB, i.e. enrichment of GPLs and concomitant depletion of lyso-GPLs in viable tumor (**Fig. 2D**). Retrospective transcript expression profiling of Lands cycle enzymes revealed overexpression of various acyltransferase genes (LPCAT1, AGPAT1, LPCAT3, MBOAT7) in GB compared to normal brain tissue but less changes in phospholipase A2 (PLA2G6) expression that underline the CPPM-based assessment. Taken together, spatial probabilistic mapping of molecular ensembles supports the global interrogation of metabolic pathways, hence opening up new avenues for the comprehensive analysis of metabolite classes.

With this study, we make the *moleculaR* framework available for the scientific community as an R package complementing leading MSI-bioinformatics packages ^20–22^. MoleculaR comes pre-loaded with the SwissLipids database and is capable of importing metabolite annotation results from the METASPACE platform to compute FDR-verified MPMs and CPPMs. *moleculaR* is equally applicable for ultra-high-resolution MSI like MRMS, or for MALDI-ToF MS and MALDI-QTOF MS. It could also be deployed and hosted on a centralized server and is equipped with a web-based GUI.

## ACKNOWLEDGEMENTS

We thank T. Alexandrov and S. Mamedov for providing guidance in all matters concerning METASPACE, and D. Hofmann for discussions concerning GB tissue. This work was supported by grants from the Deutsche Forschungsgemeinschaft (DFG, German Research Foundation) – Project-ID 404521405, SFB 1389 – UNITE Glioblastoma to T.K., A.v.D., M.P., C.A.O., W.W. and C.H., by the BMBF (German Federal Ministry of Research) as part of the Forschungscampus “Mannheim Molecular Intervention Environment” (M^2^OLIE), projects M^2^oBiTE (grant 13GW0091B) and M^2^OTAN (grant 13GW0388B) to C.H., as part of the Innovation Partnership “Multimodal Analytics and Intelligent Sensorics in Health Industry” (M^2^Aind), project M^2^OGA (grant 03FH8I02IA) to I.W. and C.H., within the framework FH-Impuls, by German Cancer Aid (70113515; Regulation of tumor immunity through the integrated stress response (ISR) in myeloid cells), and by the Klaus-Tschira Foundation, project MALDISTAR (grant 00.010.2019), to T.B. and C.H.. Acquisition of the solarix 7T XR was supported by DFG (Project 262133997) to CH.

## AUTHOR CONTRIBUTIONS

D.A.S. conceived, formulated, developed and implemented all new computational workflows, implemented the *moleculaR* R package, analyzed all MSI data, interpreted data, generated all Figures and contributed to writing of the manuscript; J.L.C. implemented the GUI and contributed to the *moleculaR* R package; J.C. implemented multimodal image fusion; C.R.G. and C.M. generated and analyzed MSI data; T.B. and I.W contributed to the theoretical formulation of the method; T.K. and W.W. planned the study, contributed GB tissue, provided clinical data and clinical tissue evaluation; M.F. and M.P. contributed GB tissue and provided clinical evaluation, A.v.D provided histopathological annotation of tissue samples; V.P., A.S. and C.A.O. performed transcript expression analysis, generated figures and contributed data interpretation; C.H. designed and coordinated the overall work, conceived applications of the computational framework, interpreted data and wrote manuscript with feedback from all co-authors.

## Methods

### Materials

All reagents were of HPLC grade. Milli-Q water (ddH2O; Millipore) was prepared in-house. Conductive indium tin oxide (ITO) coated glass slides were purchased from Bruker Daltonics (Bremen, Germany). Adhesive slides SuperFrost Plus™ were obtained from Thermo Fisher Scientific (Waltham, Massachusetts, USA) for histological analysis. Trifluoroacetic acid (TFA) and 1,5-diaminonaphtalene (1,5-DAN, ≥ 97.0%) MALDI matrix were purchased from Sigma-Aldrich (St. Louis, MO). Acetonitrile (ACN) was obtained from VWR (Darmstadt, Germany). For external calibration of the Bruker solarix magnetic resonance mass spectrometer (MRMS), a mixture of poly-DL-alanine (10 mg/mL), L-alanine ≥ 99.5% (5 mg/mL) and taurine ≥ 99% (5 mg/mL; all from Sigma Aldrich) in water was used.

### Human tissue specimen

All patients have been treated at the Heidelberg University Hospital. Patients gave informed consent prior to inclusion to exploratory molecular analysis including but not limited to MALDI Mass Spectrometry Imaging. The research is conducted in concordance with the declaration of Helsinki and was approved by the Ethics Committee at the University of Heidelberg, Germany (applications 206/2005 and AFmu-207/2017). Tissue samples were taken through primary operation of the brain tumor. Frozen resected tumor material was retrieved from the Department of Neuropathology in Heidelberg and reviewed by a board-certified neuro-pathologist. Diagnoses were molecularly confirmed according to the recent WHO classification and methylation profiles were confirmed with methylation EPIC array (#WG-317-1003, Illumina, San Diego, California, USA). Hematoxylin and eosin (H&E) stained tissues were scanned using an Aperio ImageScope scanner (Leica Biosystems) and annotated by an expert neuropathologist.

### MALDI Mass Spectrometry Imaging

Frozen human tissue was cryosectioned (10 μm; Leica CM1950; Leica Biosystems, Nussloch, Germany). Sections were mounted onto ITO slides (Bruker Daltonics), and adjacent sections were placed on SuperFrost slides (Thermo Fisher Scientific Inc., Waltham, Massachusetts, USA) for H&E staining. Cryosections were dried for 15 min in a desiccator and stored at −80 °C. Tissue sections on ITO slides were coated with 10 mg/mL 1,5-DAN matrix in 50% ACN/water using an M3 TM-Sprayer (HTX-Technologies, LLC, North Carolina, USA): Temperature: 75 °C; No of Passes: 17; Flow Rate: 0.1 mL/min; Velocity: 1200 mm/min; Track spacing: 3 mm; Pattern: CC; Pressure: 10 psi; Gas Flow Rate: 2 L/min; Nozzle Hight: 40 mm; Drying Time: 0 sec. MALDI-High-mass-resolution-imaging was performed on a solariX 7T XR (Bruker Daltonics) FTICR MRMS using Compass ftmsControl (Version 2.2.) and flexImaging (Version 5.0) softwares (both Bruker Daltonics). All measurements were done at 50 μm lateral resolution in the mass range between *m/z* 100-1200 in negative mode followed by measurement in positive mode on the same spots. Spectra were recorded with a 1M data point transient, a mass resolution of 85k at *m/z* 400, 98k at *m/z* 314, 123k at *m/z* 249 and a FID of 0.4893 sec. Per pixel 1 scan from 100 laser shots with a frequency of 1000 Hz was used. Q1 mass was set to *m/z* 120, while Time-of-Flight was adjusted to 0.9 ms. In both modes a Plate Offset of 100 V was used in combination with a deflector plate voltage of 200 V. External mass calibration was performed using polyalanine with addition of taurine (*m/z* 125.014664) to cover the whole mass range ^23^. For internal lock mass calibration in negative and positive modes, the [M-H]-signal of phosphatidylinositol (38:4) (*m/z* 885.548756) and the [M+H]+ signal of phosphatidylcholine (34:1) (*m/z* 760.585082) were used, respectively. To minimize data load, data was saved as Profile Spectrum with a Data Reduction Factor of 97%. Further data preparation and analysis were performed with in-house developed software tools as follows.

### Semi-automatic multi-modal image fusion using deformable registration

To transform H&E annotations to the MSI image domain, the optical images (5 μm^2^ per pixel) acquired prior to MALDI MSI acquisitions (and intrinsically registered with the MSI image information) and the H&E images (0.5 μm2 per pixel) were used. Briefly, H&E and optical images were transformed into grayscale images using the luminosity method (weighted average of the red, green, blue channel). Then, acquisition regions within the MSI data and annotations of the H&E files were used to define a minimal bounding box for each sample region. Subsequently, all sample regions were cropped out of the grayscale images. Cropped images of both modalities were resampled with a resolution of 7.5 μm2 per pixel. Afterwards, images were exported in the nrrd image file format. Image-based registration is used to transform the cropped images from the MSI to the H&E image domain by using elastix ^24^. The full registration is composed of a rigid step, followed by a deformable step. Each registration results in a set of parameters describing the transformation from the H&E to the MSI image domain. Those parameters are used to transform point information accordingly. Transformed polygons and corresponding annotation labels were written to mis-files.

Rigid registration is based on a multi-resolution registration strategy (Gaussian pyramid with three levels and down-sampling factors of 4,2,1). The Advanced Mattes Mutual Information metric in elastix was used as metric for the optimization of a rigid transformation using linear interpolation and 250 iterations. For the subsequent deformable registration steps, the same multi-resolution scheme and metric were applied. As deformable transformation, a recursive B-Spline transformation was used with interpolation using third-order B-Splines. The optimization was run for 750 iterations. For cases of failed image registration, a multi-metric registration approach was used, and manually defined control points were added at corresponding locations in both modalities to support the deformable registration. In this case the multi-metric output was a composition of the Matt’s Mutual Information metric and the Corresponding Points Euclidean Distance metric (equally weighted). The M^2^aia (RRID:SCR_019324; https://www.github.com/jtfcordes/m2aia) ^25^ desktop application was used to view registration results, to control registration parameters and to interactively define pairwise corresponding control points within both image modalities.

### Data Preprocessing

The centroided MALDI FTICR dataset was first exported into imzML format using SCiLS Lab Software version 2016a (Bruker Daltonics, Bremen, Germany). Further analysis proceeded in R, using the MALDIquantForeign R package for data import ^22^. Positive and negative mode spectra were stored internally in sparse-matrix representation (Matrix package) for computation efficiency. Bulk data analysis was carried out via MALDIquant ^22^. One pixel representing a full mass spectrum was randomly chosen, and full width at half maximum (fwhm) values were computed per peak and plotted against *m/z*. A smoothing curve (Friedman’s ‘super smoother’: Friedman, J. H. (1984) A variable span scatterplot smoother. Laboratory for Computational Statistics, Stanford University Technical Report No.5) was fitted to describe fwhm as a continuous function of *m/z* (**Fig. 1A**, **Suppl. Fig. 1B**) which is then used to estimate fwhm at any given *m/z*. Peaks that occurred in less than 1% of the pixels were filtered out after being binned to a relative tolerance of 12 ppm (i.e. the maximal relative deviation of a peak position to be considered as identical). Processed datasets (negative and positive modes) were exported into processed (centroided) imzML ^26^ files via MALDIquantForeign. Data-adaptive pixel-wise recalibration based on endogenous biological signals was conducted using the MSI-recalibration tool ^27^. The centroided imzML data was uploaded into the METASPACE annotation platform ^10^ and lipid search was performed against the SwissLipid database ^13^. The corresponding annotations were then downloaded as csv files and used as metabolites-of-interest (MOIs) for the molecular probability map (MPM) and collective projection map (CPPM) workflows, in other words, only METASPACE-verified MOIs were considered for subsequent analysis.

### Spatial Point Pattern (SPP) Data Representation

Given a particular MOI with its chemical sum formula and considering possible ionization states (H^-^, H^+^, K^+^ and Na^+^ for this study), the theoretical monoisotopic *m/z, m_MOI_*, is computed. This *m_MOI_* is then plugged into above mentioned fwhm continuous function to compute the expected data-dependent fwhm, which is then used to determine the *σ* of a Gaussian envelop centered at *m_MOI_* (**Fig. 1A**). This Gaussian envelope, scaled to [0,1] intensity, is used as a weighting function for any observed peaks-of-interest (POIs) occurring within its effective support (*m_MOI_* ± 3*σ*), i.e. computing 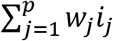 where *p* is the number of peaks observed within *m_MOI_* ± 3*σ*, and *w_j_* is the corresponding Gaussian weight at the *j*-th peak with intensity *i_j_* (**Fig. 1A** and **Suppl. Fig. 1B**). This serves as a protection against possible proximal background signals by down-weighting them relative to *m_MOI_*; the more the measured peak signal deviates from *m_MOI_*, the lower the weight it receives in the SPP representation. It is important to note, however, that this does not protect against mis-calibrated/misaligned data and in such cases we recommend applying the data-adaptive MSI recalibration method of La Rocca et al. ^27^ which was also performed for this study. The Spatstat framework ^28^ is then used to construct a marked (i.e., intensity-weighted) SPP representation *SPP_MOI_* of the MSI peaks distributed in a spatial 2D contour Φ*_tissue_* representing the tissue section with a spatial point density Λ which equals to the number of points per unit area, i.e., the average spatial density of all points *n* within Φ*_tissue_* or *n*/*A_tissue_* where *A_tissue_* is the total area of Φ*_tissue_*.

### Molecular Probability Map (MPM)

The SPP representation of MOI, *SPP_MOI_*, within a given tissue contour Φ*_tissue_* enables computation of the corresponding *MPM_MOI_*. First, a random point pattern *CSR_MOI_* is created according to a complete spatial randomness (CSR) model and is used to represent a sample of random events to be considered as an intrinsic control for every analyte case. This CSR process is generated spatially as a uniform Poisson process with a fixed spatial point density of Λ. Unlike in common CSR generating models ^8,29^, in the case of MSI, *CSR_MOI_* must also carry intensity weights (representing pixel-wise signal intensities) in order to be a valid intrinsic control model for *SPP_MOI_*. For this reason *CSR_MOI_* is then marked by signal intensities randomly sampled from the empirical intensity distribution underlying the *SPP_MOI_*. For simplicity, random sampling is replaced by randomly permuting the *SPP_MOI_* intensities, which basically has the effect of spatial reshuffling of *SPP_MOI_* points until they assume a spatial uniform Poisson process effectively dissolving any spatial clustering or spatial autocorrelation of signals (**Fig.1B**). Afterwards Kernel density estimation (KDE) is applied with an isotropic Gaussian kernel (weighted by points’ signal intensities) for both *SPP_MOI_* and its corresponding *CSR_MOI_* and is sum-normalized to compute the spatial density functions *ρ_MOI_*(*x, y*) and *ρ_CSR_*(*x, y*), respectively. Let *f_MOI_*(*k*) and *f_CSR_*(*k*) denote the probability density functions of intensities *k* obtained from the resulting *ρ_MOI_*(*x, y*) and *ρ_CSR_*(*x,y*), respectively. As a consequence of the central limit theorem, the intensity distribution *f_CSR_*(*k*) converges towards a normal distribution as the bandwidth increases to infinity, which in practice can already be observed for small bandwidth values. This does not necessarily apply to *f_MOI_*(*k*) (see **Suppl. Fig. 3a** and **b**).

Hence

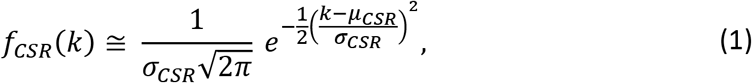

where *μ_CSR_* and *σ_CSR_* are the mean and standard deviation of *ρ_CSR_*(*x,y*). To identify areas which have a higher likelihood of showing a significant abundance of MOI when compared to a random distribution (i.e. analyte hotspots) and, on the other hand, areas which have a higher likelihood of showing a significant deficiency of MOI (i.e. analyte coldspots), upper and lower cutoff quantiles *k_upr_* and *k_lwr_* are defined for *f_CSR_*(*k*), respectively:

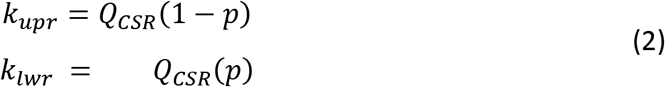

Here, *Q_CSR_*(*p*) denotes the corresponding quantile function (i.e. inverse cumulative distribution function) of *f_CSR_*, and *p* is the p-value threshold below which the null hypothesis is rejected (set to 0.05 for this study). To account for the inherent multiple testing problem, a conservative Bonferroni correction (used for all figures in this study) or, optionally, a less conservative Benjamini-Hochberg correction is applied. The analyte hotspots and coldspots are accordingly defined as locations (*x_hs_, y_hs_*) and (*x_cs_, y_cs_*) where

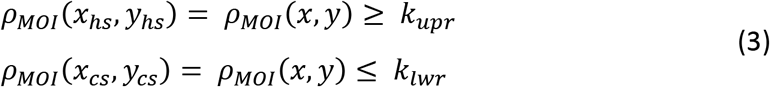

The MOI’s molecular probability map, *MPM_MOI_* is then defined as a composite representation of MOI spatial density of Gaussian weighted intensities according to the scheme shown in **Fig. 1A**, with analyte hotspots and/or analyte coldspots superimposed as polygonal contours identifying areas of MOI significant abundance and deficiency, respectively. The bandwidth *h_G_* for the chosen Gaussian kernel of the KDE is computed in two distinct ways; KDE is applied iteratively with *h_G_* varying from 1 to 10 (pixels; multiples of 50 μm in this study) in 0.5 increments, during each iteration the Moran’s I statistic, a measure of spatial autocorrelation, is determined. The optimal *h_G_* is then determined by finding the point in the Moran’s I vs *h_G_* plot at which the spatial autocorrelation levels-off, i.e. after which an increase in *h_G_* does not result in a considerable increase in the spatial autocorrelation of the smoothed density image (**Suppl. Fig. 2**). This “elbow” point is determined by finding the maximum distance from points on the curve to a line drawn between the curve’s end points (**Suppl. Fig. 2**). For CPPM, *h_G_* is more efficiently computed by the Scott method i.e. *h_G_* = *n*^−1/(*d*+4)^; where d is the number of dimensions and, for this study, equals to 2 ^30^.

### Collective Projections Probability Map (CPPM)

Given a set of MOIs *C* ∈ {*MOI*_1_, *MOI*_2_, …, *MOI_m_*}*x*, for each single MOI_i_ an SPP representation *SPP_i_* is calculated as previously described. Afterwards all individual *SPP_i_* are projected into the same Φ*_tissue_* allowing for coordinate duplication resulting in *SPP_C_* where the overlap of points results in accumulation of the corresponding signal intensities. The workflow then commences with the square root transformation of intensities to compensate for the inherent heteroskedasticity and possible differences in ionization efficiency between the individual MOI_i_s. Then *CSR_C_* is created and subsequently KDE is applied on both sides until *MPM_C_* is computed as described in the previous section such that the resulting collective projection probability map *CPPM_C_* is equivalent to *MPM_C_*. The naming distinction is only made to emphasize that CPPM is based on the visualization of multiple analytes at a time.

For any number of MOIs, basic arithmetic operations on the spatial point patterns of MOIs could also be applied. This is useful when a ratio of two MOIs is desired (**ex. Fig. 2B** bottom row) or when a more complex evaluation is of interest (**ex. Fig. 2B** top right and bottom right). To perform such operations, first the set of the input *SPP_MOI_*s are converted into pixel-based images with equal pixel grids. Afterwards the spatial expression is evaluated on a pixel-by-pixel basis. Possible divisions by zero are computationally dropped. The resulting raster image is then converted back to an SPP whose points are carrying the respective computed pixel intensities. The created SPP is then fed into the MPM framework as previously described.

### Transcript Expression Profiling of TCGA and GTEx Datasets

TCGAbiolinks ^31^ was used to download fragments per kilobase of transcript per million mapped reads (FPKM) and the clinical information of The Cancer Genome Atlas (TCGA) glioblastoma (GB) datasets from Genomic Data Commons (GDC) (https://gdc.cancer.gov). Patient samples characterized as “primary tumor” were retained (n = 156). The FPKM values were converted to transcripts per million (TPMs) ^32^. TPM data of normal brain tissues (n = 1671) were downloaded from the Genotype-Tissue Expression (GTEx) dataset (https://gtexportal.org). All TPM values were log2-transformed.

For bioinformatics analysis of TCGA and GTEx data, all pairwise comparisons were performed using Kruskal-Wallis and Wilcoxon rank-sum tests. All analyses were run in R (https://cran.r-project.org) version 4.1, and Bioconductor (https://bioconductor.org) version 3.14. All graphical representations were generated using ggplot2, RColorBrewer, gridExtra, and ggridges packages.

### Data Availability

Raw data that support the anticipated results is available at Metaspace through the following link: https://metaspace2020.eu/project/abusammour-2021.

### Code Availability

A well-documented companion R package that implements the presented framework is made available at https://github.com/CeMOS-Mannheim/moleculaR alongside with clear introductory sections and exemplary code vignettes. The R package is equipped with a web-based GUI and could be deployed and hosted on a centralized server as described in package link above.

## REFERENCES

1 Schulz, S., Becker, M., Groseclose, M. R., Schadt, S. & Hopf, C. Advanced MALDI mass spectrometry imaging in pharmaceutical research and drug development. Curr Opin Biotechnol 55, 51–59, doi:10.1016/j.copbio.2018.08.003 (2019).

2 Scupakova, K. et al. Cellular resolution in clinical MALDI mass spectrometry imaging: the latest advancements and current challenges. Clin Chem Lab Med 58, 914–929, doi:10.1515/cclm-2019-0858 (2020).

3 Van de Plas, R., Yang, J., Spraggins, J. & Caprioli, R. M. Image fusion of mass spectrometry and microscopy: a multimodality paradigm for molecular tissue mapping. Nat Methods 12, 366–372, doi:10.1038/nmeth.3296 (2015).

4 Niehaus, M., Soltwisch, J., Belov, M. E. & Dreisewerd, K. Transmission-mode MALDI-2 mass spectrometry imaging of cells and tissues at subcellular resolution. Nat Methods 16, 925–931, doi:10.1038/s41592-019-0536-2 (2019).

5 Balluff, B., Hopf, C., Porta Siegel, T., Grabsch, H. I. & Heeren, R. M. A. Batch Effects in MALDI Mass Spectrometry Imaging. J Am Soc Mass Spectrom 32, 628–635, doi:10.1021/jasms.0c00393 (2021).

6 Race, A. M. & Bunch, J. Optimisation of colour schemes to accurately display mass spectrometry imaging data based on human colour perception. Anal Bioanal Chem 407, 2047–2054, doi:10.1007/s00216-014-8404-5 (2015).

7 Lee, J., Narang, S., Martinez, J., Rao, G. & Rao, A. Spatial Habitat Features Derived from Multiparametric Magnetic Resonance Imaging Data Are Associated with Molecular Subtype and 12-Month Survival Status in Glioblastoma Multiforme. PLoS One 10, e0136557, doi:10.1371/journal.pone.0136557 (2015).

8 Kather, J. N. et al. Continuous representation of tumor microvessel density and detection of angiogenic hotspots in histological whole-slide images. Oncotarget 6, 19163–19176, doi:10.18632/oncotarget.4383 (2015).

9 Peters, R. et al. Quantification of fibrous spatial point patterns from single-molecule localization microscopy (SMLM) data. Bioinformatics 33, 1703–1711, doi:10.1093/bioinformatics/btx026 (2017).

10 Palmer, A. et al. FDR-controlled metabolite annotation for high-resolution imaging mass spectrometry. Nat Methods 14, 57–60, doi:10.1038/nmeth.4072 (2017).

11 Cassese, A. et al. Spatial Autocorrelation in Mass Spectrometry Imaging. Anal Chem 88, 5871–5878, doi:10.1021/acs.analchem.6b00672 (2016).

12 Friedrich, M. et al. Tryptophan metabolism drives dynamic immunosuppressive myeloid states in IDH-mutant gliomas. Nature Cancer 2, 723–740, doi:https://doi.org/10.1038/s43018-021-00201-z (2021).

13 Aimo, L. et al. The SwissLipids knowledgebase for lipid biology. Bioinformatics 31, 2860–2866, doi:10.1093/bioinformatics/btv285 (2015).

14 Garate, J. et al. A Drastic Shift in Lipid Adducts in Colon Cancer Detected by MALDI-IMS Exposes Alterations in Specific K(+) Channels. Cancers (Basel) 13, doi:10.3390/cancers13061350 (2021).

15 Fernandez, R. et al. Identification of Biomarkers of Necrosis in Xenografts Using Imaging Mass Spectrometry. J Am Soc Mass Spectrom 27, 244–254, doi:10.1007/s13361-015-1268-x (2016).

16 Atkinson, D. E. & Walton, G. M. Adenosine triphosphate conservation in metabolic regulation. Rat liver citrate cleavage enzyme. The Journal of Biological Chemistry 242, 3239–3241 (1967).

17 Hayakawa, E., Fujimura, Y. & Miura, D. MSIdV: a versatile tool to visualize biological indices from mass spectrometry imaging data. Bioinformatics 32.24, 3852–3854 (2016).

18 Deen, S. S. et al. Sodium MRI with 3D-cones as a measure of tumour cellularity in high grade serous ovarian cancer. Eur J Radiol Open 6, 156–162, doi:10.1016/j.ejro.2019.04.001 (2019).

19 Guo, S., Wang, Y., Zhou, D. & Li, Z. Significantly increased monounsaturated lipids relative to polyunsaturated lipids in six types of cancer microenvironment are observed by mass spectrometry imaging. Sci Rep 4, 5959, doi:10.1038/srep05959 (2014).

20 Bemis, K. D. et al. Cardinal: an R package for statistical analysis of mass spectrometry-based imaging experiments. Bioinformatics 31, 2418–2420, doi:10.1093/bioinformatics/btv146 (2015).

21 Rafols, P. et al. rMSI: an R package for MS imaging data handling and visualization. Bioinformatics 33, 2427–2428, doi:10.1093/bioinformatics/btx182 (2017).

22 Gibb, S. & Strimmer, K. MALDIquant: a versatile R package for the analysis of mass spectrometry data. Bioinformatics 28, 2270–2271, doi:10.1093/bioinformatics/bts447 (2012).

## References

23 Gruendling, T., Sauerland, V., Barahona, C., Herz, C. & Nitsch, U. Polyalanine - a practical polypeptide mass calibration standard for matrix-assisted laser desorption/ionization mass spectrometry and tandem mass spectrometry in positive and negative mode. Rapid Commun Mass Spectrom 30, 681–683, doi:10.1002/rcm.7492 (2016).

24 Klein, S., Staring, M., Murphy, K., Viergever, M. A. & Pluim, J. P. elastix: a toolbox for intensity-based medical image registration. IEEE Trans Med Imaging 29, 196–205, doi:10.1109/TMI.2009.2035616 (2010).

25 Cordes, J. et al. M2aia-Interactive, fast, and memory-efficient analysis of 2D and 3D multi-modal mass spectrometry imaging data. Gigascience 10, giab049, doi:10.1093/gigascience/giab049 (2021).

26 Schramm, T. et al. imzML--a common data format for the flexible exchange and processing of mass spectrometry imaging data. J Proteomics 75, 5106–5110, doi:10.1016/j.jprot.2012.07.026 (2012).

27 La Rocca, R. et al. Adaptive Pixel Mass Recalibration for Mass Spectrometry Imaging Based on Locally Endogenous Biological Signals. Anal Chem 93, 4066–4074, doi:10.1021/acs.analchem.0c05071 (2021).

28 Baddeley, A., Rubak, E. & Turner, R. Spatial Point Patterns - Methodology and Applications with R. (Chapman and Hall/CRC, 2016).

29 Kuhldorff, M. A spatial scan statistic. Communications in Statistics - Theory and Methods 26, 1481–1496, doi:doi.org/10.1080/03610929708831995 (1997).

30 Scott, D. W. Multivariate Density Estimation: Theory, Practice, and Visualization. (John Wiley & Sons, Inc., 1992).

31 Colaprico, A. et al. TCGAbiolinks: an R/Bioconductor package for integrative analysis of TCGA data. Nucleic Acids Res 44, e71, doi:10.1093/nar/gkv1507 (2016).

32 Li, B. & Dewey, C. N. RSEM: accurate transcript quantification from RNA-Seq data with or without a reference genome. BMC Bioinformatics 12, 323, doi:10.1186/1471-2105-12-323 (2011).

