## Supplementary Figures for "Spatial Probabilistic Mapping of Metabolite Ensembles in Mass Spectrometry Imaging"

**Suppl. Figure 8.** CPPMs for of an adjacent tissue section of GBM tissue section showcased in Fig. 2. . 10

### Supplementary Figures

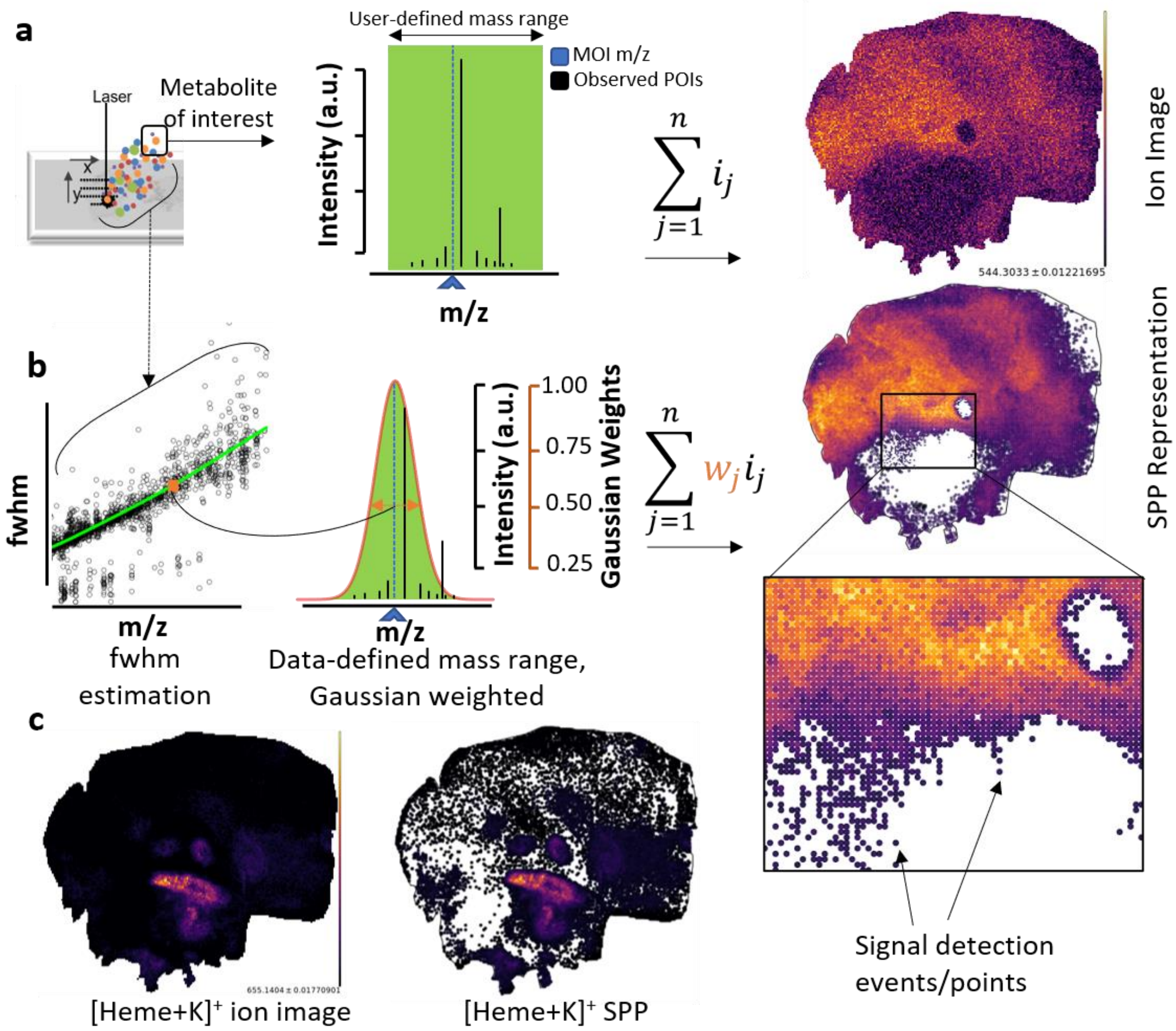

**Suppl. Figure 1. Schematic diagram illustrating the concepts of ion image versus spatial point pattern (SPP) representation of an observed Peak-of-Interest (POI) and a database-annotated Metabolite-of-interest (MOI)**, which may or may not be identical. A) Ion images are generated by summing up observed ion intensities (dark vertical lines) within a user-defined mass-range (green area) around an observed POI. MOI (dark blue arrow and dotted blue line) that users want to analyze and visualize and that may or may not correspond to POI, require statistically valid spatial mapping. B) SPP representation of the same metabolite. A user-selected full mass spectrum is used to compute full width half maximum (fwhm) values of its peaks across the m/z axis. A curve is fitted to describe the continuous relation of fwhm as a function of m/z. A Gaussian envelope, whose sigma is inferred from the fwhm model at MOI, is centered on MOI. The Gaussian is then used as a weighting factor to protect against proximal background signals; the further the measured m/z (= POI) from the theoretical m/z (= MOI), the lower the weight it receives in the SPP representation. The resulting SPP map represents weighted points/events distributed in a 2D window, i.e. the tissue section. C) An exemplary comparison between standard ion image and SPP representation of $[Heme+K]^+$.

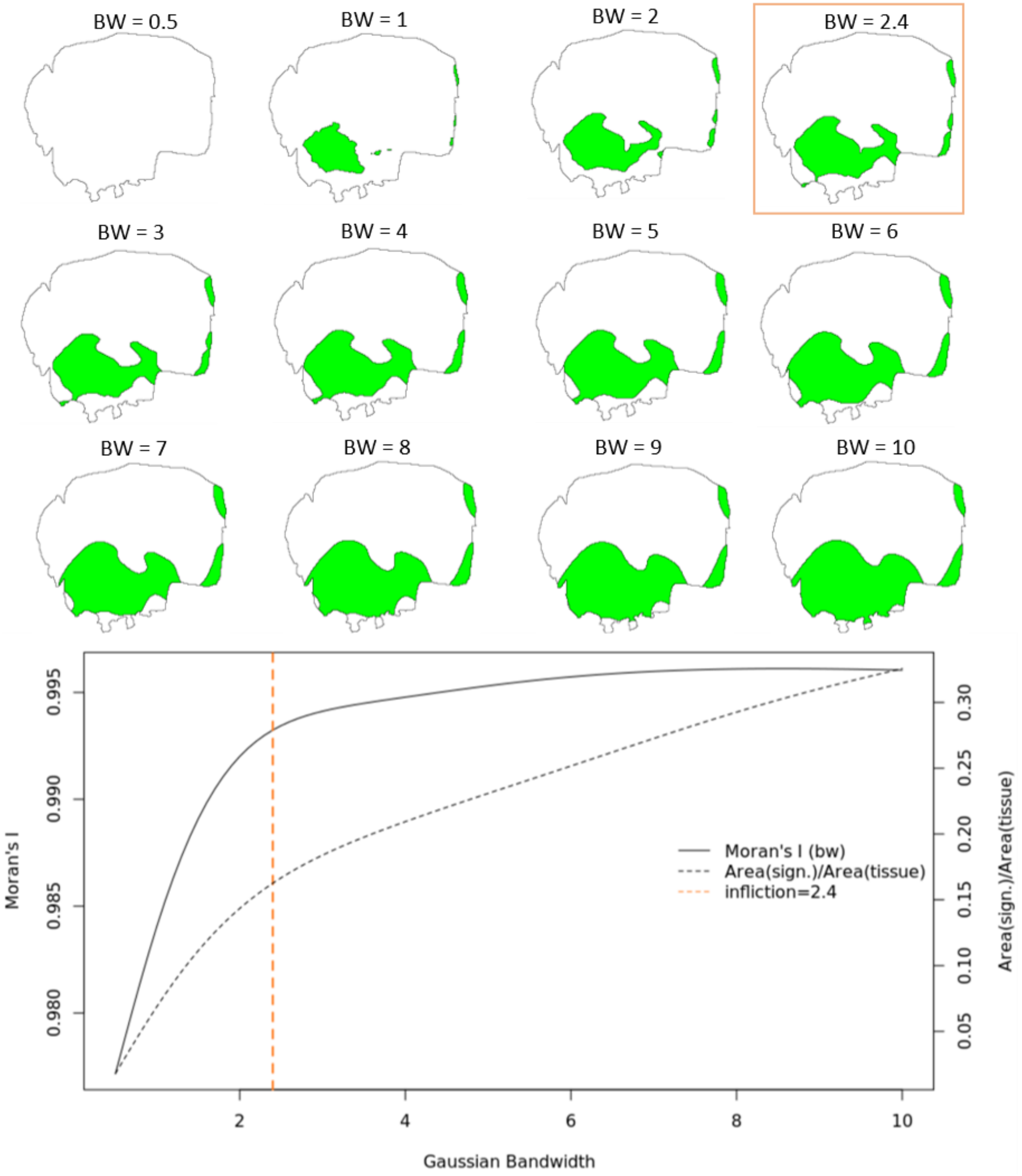

**Suppl. Figure 2. Bandwidth estimation for the chosen Gaussian kernel of the kernel density estimation procedure as part of the MPM workflow** (**Fig. 1B**) evaluated for PS(36:1)-H$^-$ at *m/z* 788.5447 (**Fig. 1E**). Bandwidth $h_G$ is varied iteratively from values of 1 to 10 (pixels; multiples of 50 µm) in 0.5 incremental steps, during each iteration KDE is applied and the Moran's I statistic, a measure of autocorrelation, is determined. The optimal $h_G$ is then determined by finding the point in the Moran's I vs $h_G$ plot at which the spatial autocorrelation levels-off, i.e. after which an increase in $h_G$ does not result in a considerable increase in the spatial autocorrelation of the smoothed density image. This "elbow" point is determined by finding the maximum distance from points on the curve to a line drawn between the curve's end points. Additionally, the dashed line shows how the statistically significant area (i.e. analyte hotspot) changes relative to the total tissue area as a function of $h_G$.

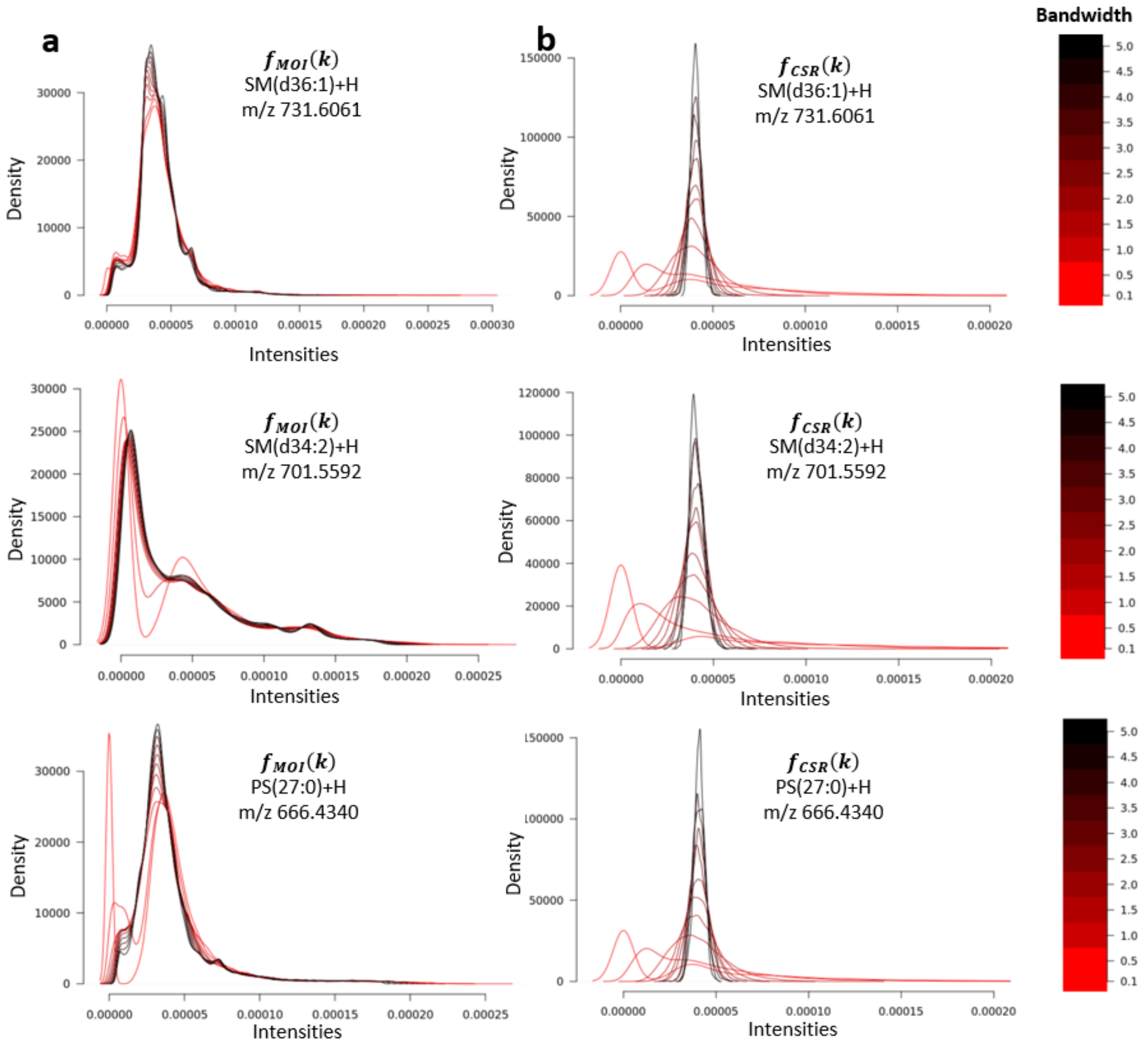

**Suppl. Figure 3. 1D Intensity distributions $f_{MOI}(k)$ (column a) and $f_{CSR}(k)$ (column b) of three different lipids (rows) corresponding to the 2D spatial densities $\rho_{MOI}(x,y)$ and $\rho_{CSR}(x,y)$, respectively.** While $f_{MOI}(k)$ does not necessarily converge to a normal distribution for the range of bandwidths under consideration (part a), as a consequence of the central limit theorem, the corresponding intensity distribution $f_{CSR}(k)$ approximates a normal distribution as the bandwidth increases, even for smaller bandwidth values (part b).

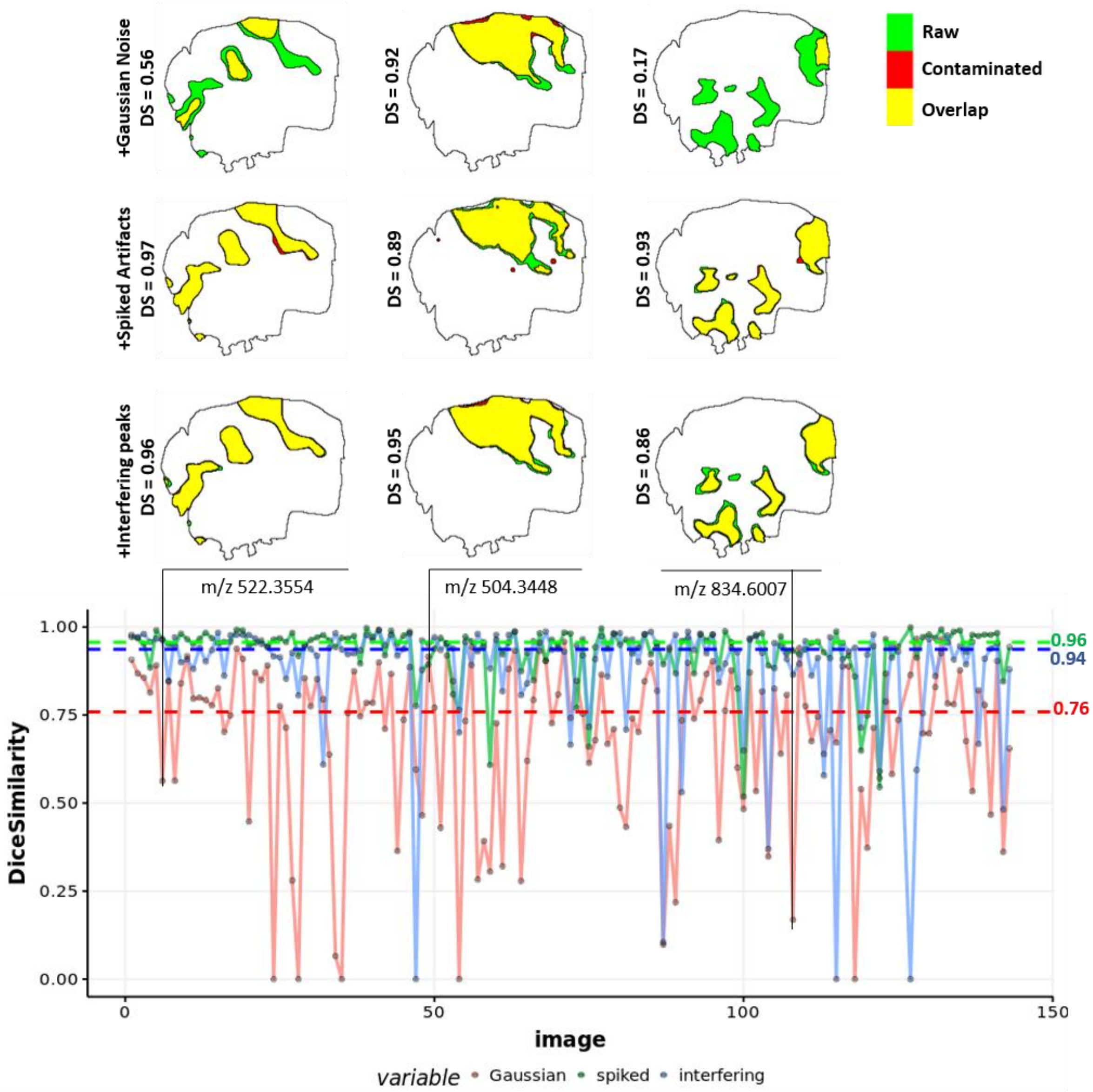

**Suppl. Figure 4. Dice similarity coefficient computed for areas of significance (analyte hotspots) between raw MSI against contaminated data of 142 MPM images of MOIs (identified in Metaspace with ≤ 0.2 FDR) in positive ion mode.** Noise contamination was based on random Gaussian noise added to POI (first row of upper panel; red curve in lower panel), presence of abnormally high-intensity peak artefacts (second row of upper panel; green curve in lower panel) and added overlapping analytes occurring within the span of the theoretical Gaussian envelop specific for MOI (2σ away from MOI m/z; third row of upper panel; blue curve in lower panel). Median Dice similarity coefficients were 0.76 (dashed red line), 0.96 (dashed green line) and 0.94 (dashed blue line) for the contamination sources Gaussian noise, spiked artefacts and interfering peaks, respectively.

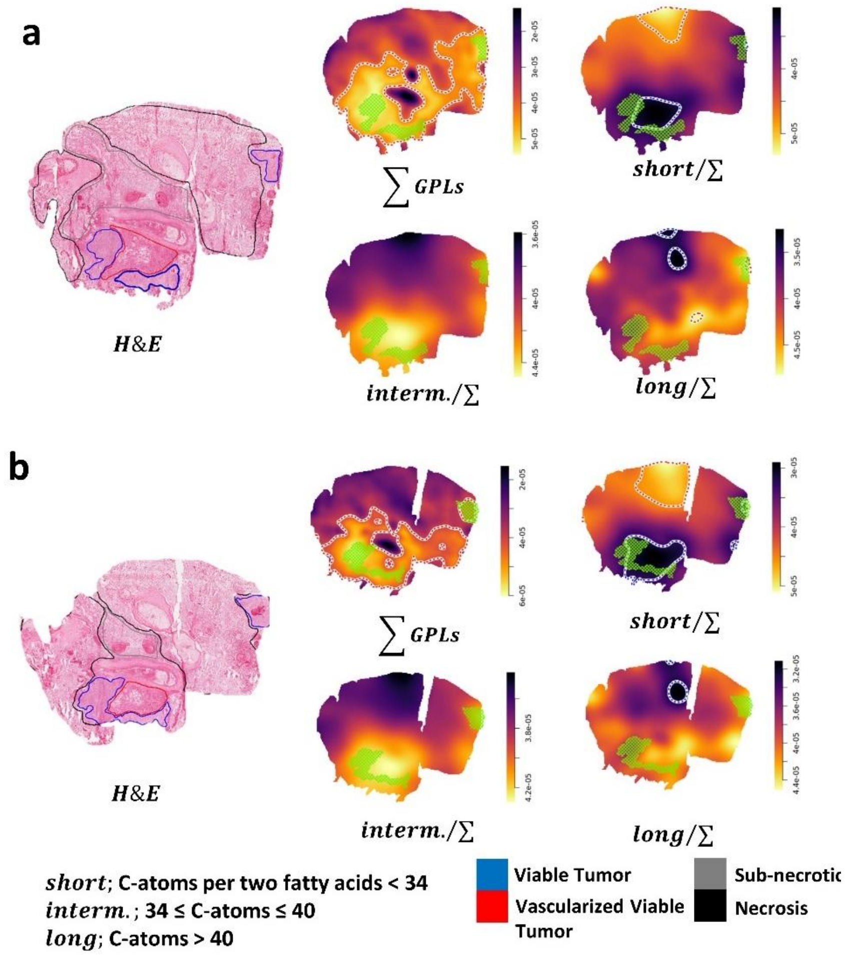

**Suppl. Figure 5. Fatty acid chain length-focused visualization for short (< 34C), intermediate ([34C,40C]) and long (> 40C) and their corresponding sum-normalized CPPMs of GPLs (i.e. $\sum GPLs$; [M+K$^+$], [M+Na$^+$] and [M+H$^+$] of PC, PE, PS, PI) for one Glioblastoma tissue section (a) and its serial tissue section (b).**

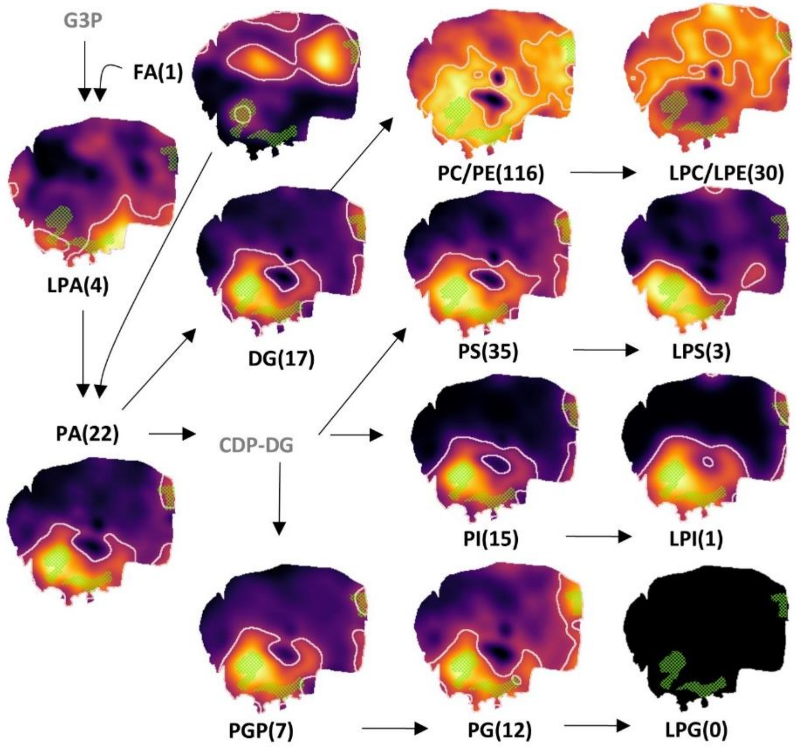

**Suppl. Figure 6. CPPMs of lipid classes of the GBM tissue section showcased in Fig. 1. CPPMs enable spatial investigation of lipid synthesis and remodelling pathways in GBM tissue sections by visualizing structurally similar analytes (e.g., lipid classes) within the same image space.** Numbers in parenthesis refer to all detected and METASPACE-identified lipids at ≤ 0.2 FDR incl. alkali adducts. FA, fatty acid; PA, phosphatidic acid; DG, diglycerides; PC, phosphatidylcholine; LPC, lysophosphatidylcholine; LPE, lysophosphatidylethanolamine; PE, phosphatidylethanolamine; PI, phosphatidylinositol; CDP-DG, cytidinediphosphatediacyl glycerol; LPG, lysophosphatidylglycerol; PG, phosphatidylglycerol; PGP, phosphatidylglycerolphosphate; LPS, lysophosphatidylserine; PS, phosphatidylserine.

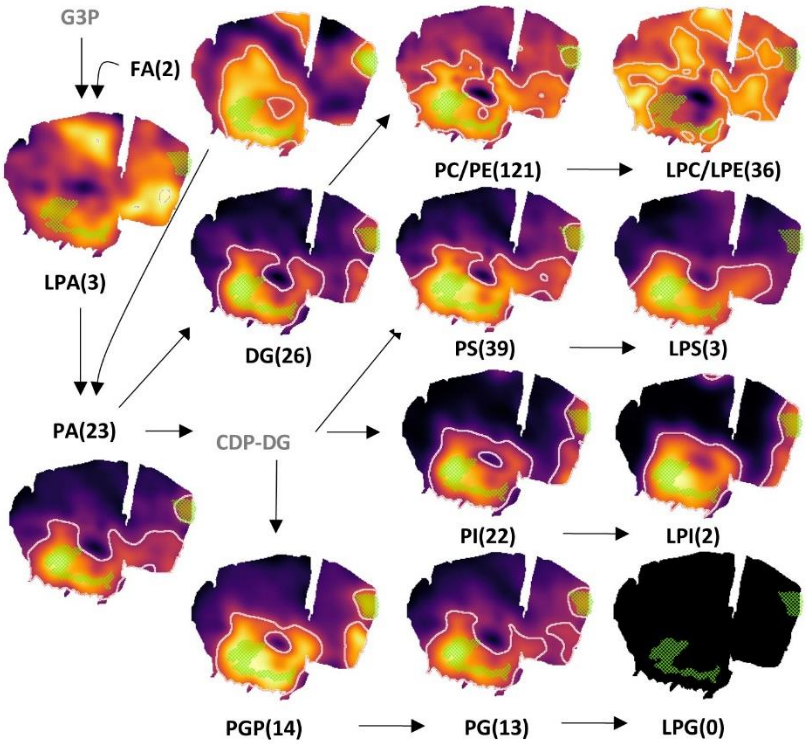

**Suppl. Figure 7. CPPMs of lipid classes of a serial tissue section of GBM tissue section showcased in Fig. 1. CPPMs enable spatial investigation of lipid synthesis and remodelling pathways in GBM tissue sections by visualizing structurally similar analytes (e.g., lipid classes) within the same image space.** Numbers in parenthesis refer to all detected and METASPACE-identified lipids at ≤ 0.2 FDR incl. alkali adducts. FA, fatty acid; PA, phosphatidic acid; DG, diglycerides; PC, phosphatidylcholine; LPC, lysophosphatidylcholine; LPE, lysophosphatidylethanolamine; PE, phosphatidylethanolamine; PI, phosphatidylinositol; CDP-DG, cytidinediphosphatediacyl glycerol; LPG, lysophosphatidylglycerol; PG, phosphatidylglycerol; PGP, phosphatidylglycerolphosphate; LPS, lysophosphatidylserine; PS, phosphatidylserine.

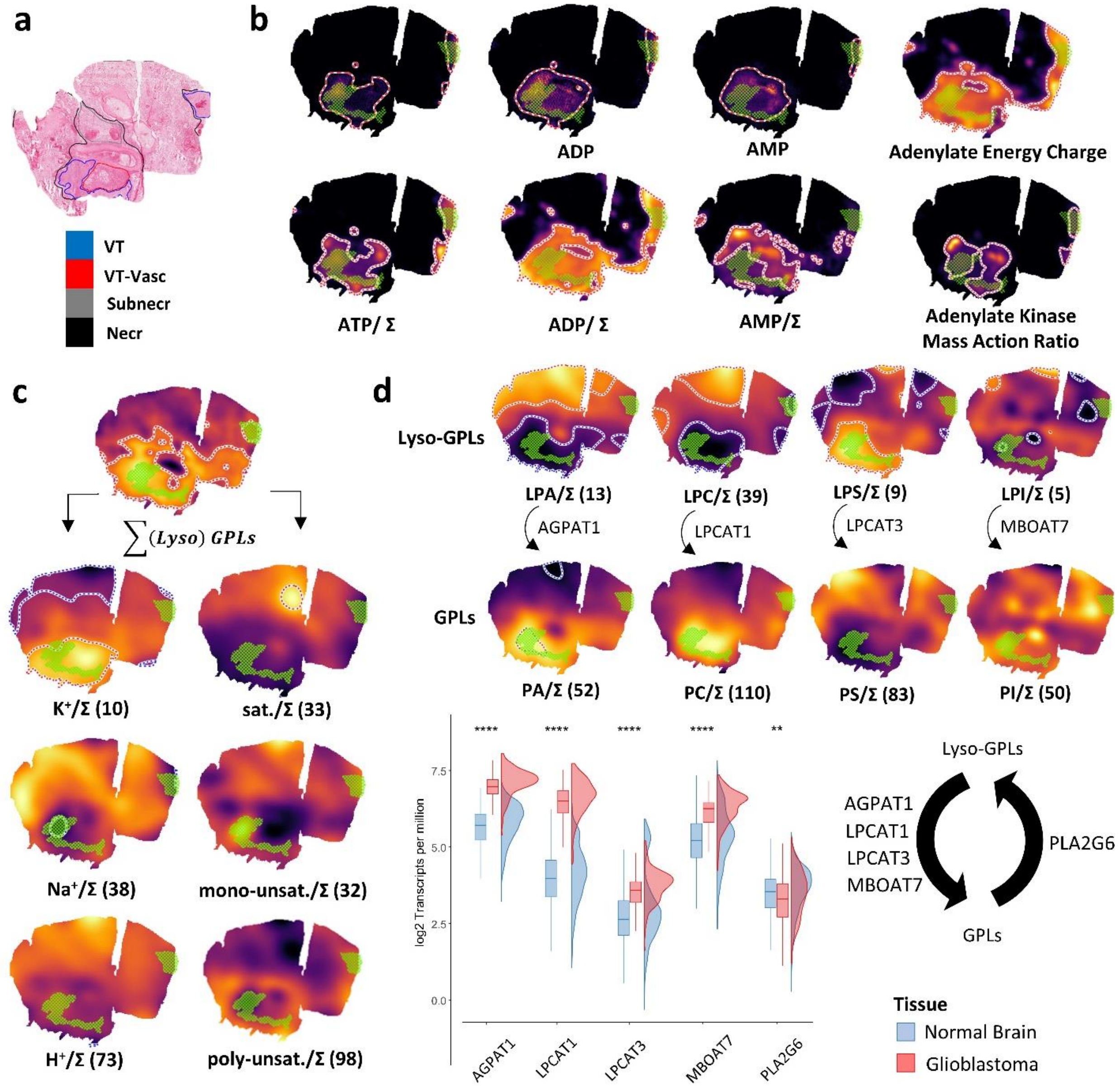

**Suppl. Figure 8. CPPMs for of an adjacent tissue section of GBM tissue section showcased in Fig. 2.** a) H&E-stained image of the tissue section highlighting different GBM anatomical regions (VT: vital tumor; VT-Vasc: vital tumor with high vascularization; Subnecr: areas of pre-necrotic tissue; Necr: necrotic tissue ). **b**) CPPMs enable basic arithmetic manipulations on SPPs of multiple MOIs. Green mesh indicates co-registered vital tumor regions (part **a**). MPMs of [ATP-H]$^-$, [ADP-H]$^-$ and [AMP-H]$^-$ (≤0.2 FDR; upper row) compared to their sum-normalized CPPMs (bottom row; Σ = [ATP-H]$^-$ + [ADP-H]$^-$ + [AMP-H]$^-$ ). CPPMs also enable complex spatial quantitative scores such as adenylate energy charge (([ATP-H]$^-$ +0.5 [ADP-H]$^-$)/([ATP-H]$^-$+[ADP-H]$^-$+[AMP-H]$^-$); top right) and adenylate kinase mass action ratio ([ATP-H]$^-$[AMP-H]$^-$/[ADP-H]$^{-2}$; bottom right). **c**) Analysis of the tissue's alkali ion milieu and lipid (un)saturation. Numbers in parenthesis = METASPACE-verified lipids at ≤ 0.2 FDR. left column: CPPMs of all detected protonated or alkali metal adducts of (lyso-)GPLs (PC, LPC, PE, LPE, PS, LPS, PI, LPI) relative to the overall sum Σ(lyso)GPLs. right column: CPPMs showing lipid (un)saturation relative to Σ(lyso)GPLs (sat: saturated; unsat: unsaturated). **d**) CPPMs enable spatial investigation of glycerolphospholipid remodeling (Lands' cycle) in GB by visualizing structurally similar lipids (≤ 0.5 FDR) within the same image space. Upper panel: Lyso- and non-lyso-GPL pairs are normalized to their sum (ex. for LPC and PC, Σ represents the sum of all LPC and PC lipids). Lower panel (same data shown as in **Fig 2D** put here for convenience): Rainfall plot showing boxplot and density curves representations of the expression of select Lands' cycle enzymes in normal brain (blue; GTEx data) and GB (red; TCGA data) both represented as log2 transcripts per million (Wilcoxon rank-sum test, ** $P < 0.01$, **** $P < 0.0001$). PA, phosphatidic acid; PC, phosphatidylcholine; LPC, lysophosphatidylcholine; LPE, lysophosphatidylethanolamine; PE, phosphatidylethanolamine; PI, phosphatidylinositol; LPG, lysophosphatidylglycerol; PG, phosphatidyl-glycerol; LPS, lysophosphatidylserine; PS, phosphatidylserine.
